# Jk DNA GAGA MOTIFS ARE REQUIRED FOR LOCAL NUCLEOSOME REMODELING AND Vk-Jk RECOMBINATION

**DOI:** 10.1101/2025.01.21.634169

**Authors:** Margaret Veselits, Kaitlin McLean, Nathaniel E Wright, Michael Okoreeh, Mark Maienschein-Cline, Malay Mandal, Marcus Clark

**Affiliations:** Department of Medicine, Section of Rheumatology, University of Chicago, Chicago, IL, 60637, USA; Gwen Knapp Center for Lupus and Immunology Research, University of Chicago, Chicago, IL, 60637, USA; Core for Research Informatics, University of Illinois at Chicago, Chicago, IL, 60612 USA

## Abstract

Immunoreceptor gene recombination requires complementary 12 bp and 23 bp recombination signal sequences (RSSs). In addition, the RSSs that assemble the RAG proteins, recombination centers, must be accessible yet flanked by a 5’ nucleosome decorated with H3K4me3. In *Drosophila*, DNA GAGA motifs play an important role in nucleosome positioning. Herein, we report that 5’ to each functional Jk RSS is a DNA GAGA motif conserved across mammalian species. In mice, the GAGA motif 5’ to Jk1 regulated local RSS accessibility and 5’ nucleosome placement. Furthermore, it was required for Vk-Jk1 recombination. Murine Jk3 is nonfunctional, having mutations in both RSS and GAGA motifs. Restoring both GAGA and RSS motifs rescued Vk-Jk3 recombination. In contrast, restoring the RSS alone did not. Genome-wide, strong cryptic 23 RSSs were preferentially bound to nucleosomes. Furthermore, evolutionary selection against cRSS only occurred in the A Compartment of B lymphocytes, not embryonic stem cells. These data indicate that in developing B cells, nucleosome positioning both enables and restricts recombination to Jk. Furthermore, our data suggest an expanded definition of recombination center-associated RSSs to include a 5’ GAGA sequence that dictates the local epigenetic state required for gene recombination.

**Summary:** Recombination center assembly requires a specific epigenetic topology at recombination signal sequences. Herein, we report that conserved GAGA motifs 5’ to each Jk segment are required for establishing this epigenetic topology and subsequent local gene recombination.

## Introduction

B lymphocytes produce a vast repertoire of antibodies to protect from a myriad of pathogens. This diversity is largely achieved through stochastic V(D)J recombination of the immunoglobulin (Ig) genes (*1, 2*). Ig gene cleavage is mediated by proteins encoded by the two recombinase activating genes, *RAG1* and *RAG2* (*3, 4*). RAG protein-induced double strand breaks (DSBs) occur at specific recombination signal sequences (RSSs) containing highly conserved nonamer and heptamer motifs, separated by 12 or 23 base pairs. The “12/23 Rule” dictates that recombination must occur between one RSS containing a 12 bp spacer and one RSS containing a 23 bp spacer (*5–7*). At *Igk*, each Vk gene segment has a 3’ 12 bp RSS and each functional Jk gene segment has a 5’ 23 bp RSS. The RAG complex assembles at Jk RSSs to form recombination centers to which the Vk gene segments are recruited (*1, 2, 8, 9*).

While gene recombination is necessary for antigen receptor diversity, any mistargeting of the RAG complex risks genomic translocations and malignant transformation (*10–12*). Indeed, throughout the genome there are cryptic RSSs (cRSSs), which can be cleaved by the RAG complex and lead to genomic instability (*13, 14*). To mitigate this risk, mechanisms have evolved to ensure the fidelity of the RAG complex.

For example, the exquisite temporal, cell cycle, and tissue-specific regulation of RAG expression restricts gene recombination to specific lymphocyte developmental states (*2–4, 15, 16*). Furthermore, RAG recruitment to DNA is regulated epigenetically as RAG2 binds nucleosomes decorated with histone H3 lysine 4 trimethylation (H3K4me3) via its plant homeodomain (PHD) finger domain (*17–19*). Thus, RAG-mediated recombination requires the local presence of nucleosomes.

Paradoxically, when bound *in vitro* to nucleosomes, RSSs are resistant to cleavage by RAG (*20, 21*). Furthermore, nucleosomes preferentially bind RSS-containing DNA sequences (*22*). These data indicate that nucleosomes bearing H3K4me3 required for RAG2 recruitment must be precisely positioned at Jk to both recruit RAG2 yet allow RAG1-mediated RSS cleavage (*2*). Indeed, this necessary topology is enforced by the epigenetic reader BRWD1, which remodels local nucleosome structure to both place a nucleosome 5’ to each Jk RSS and to ensure each RSS is accessible (*23*).

Genome-wide, there is a striking association between BRWD1-dependent nucleosome remodeling and the presence of GA-repeats in DNA (“GAGA motifs”) (*23*). In *Drosophila*, the epigenetic modifier GAGA Factor (GAF) plays a critical role in the expression of homeotic genes (*24, 25*). GAF is a pioneer transcription factor that binds to GA-rich DNA sequences, with a GAGAG pentamer being the canonical consensus sequence (*26*). However, GAF can bind a GAG trinucleotide at minimum (*27*). GAF works with chromatin remodeling complexes in an ATP-dependent manner to slide or evict nucleosomes located in proximity to GAGA motifs (*28–30*). However, the role of GAGA motifs in modulating nucleosome positioning in mammals is unknown.

Herein, we report that conserved GAGA motifs located 5’ to each Jk RSS play a crucial role in establishing the nucleosome architecture required for efficient *Igk* recombination. Deletion and add-back experiments demonstrate that GAGA motifs are both necessary and sufficient to dictate nucleosome structure around Jk 23 bp RSSs and enable Vk-Jk recombination in small pre-B cells. Furthermore, cryptic 23 bp RSSs (cRSSs) genome-wide were preferentially occupied by nucleosomes. Taken together, these data suggest that nucleosome positioning is a critical component of the epigenetic regulation that both enables and restricts cleavage to canonical RSSs in B lymphocytes.

## Results

### The *Jk1 5’ GAGA region is required for Igk recombination*

Examination of nucleotide sequences at the Jk locus revealed that each functional Jk gene segment was preceded by a GAGAG DNA sequence (“GAGA motif”) located an average of approximately 80 nucleotide base pairs (bp) 5’ to each RSS (**Figure 1A**). Interestingly, the non-functional Jk3 gene segment was not associated with a GAGAG motif. Furthermore, a comparison of the sequence 5’ to the *Igk* Jk1 segment across various vertebrate species indicated that this motif was conserved (**Supplementary Figure 1A**). In most species examined, it was located within 80 bp of the Jk1 RSS, while in others it was located approximately 350 bp upstream. We hypothesized that if this motif is important for nucleosome positioning and *Igk* recombination, its removal should result in reduced usage of the corresponding Jk in the expressed *Igk* repertoire.

**Figure 1.**
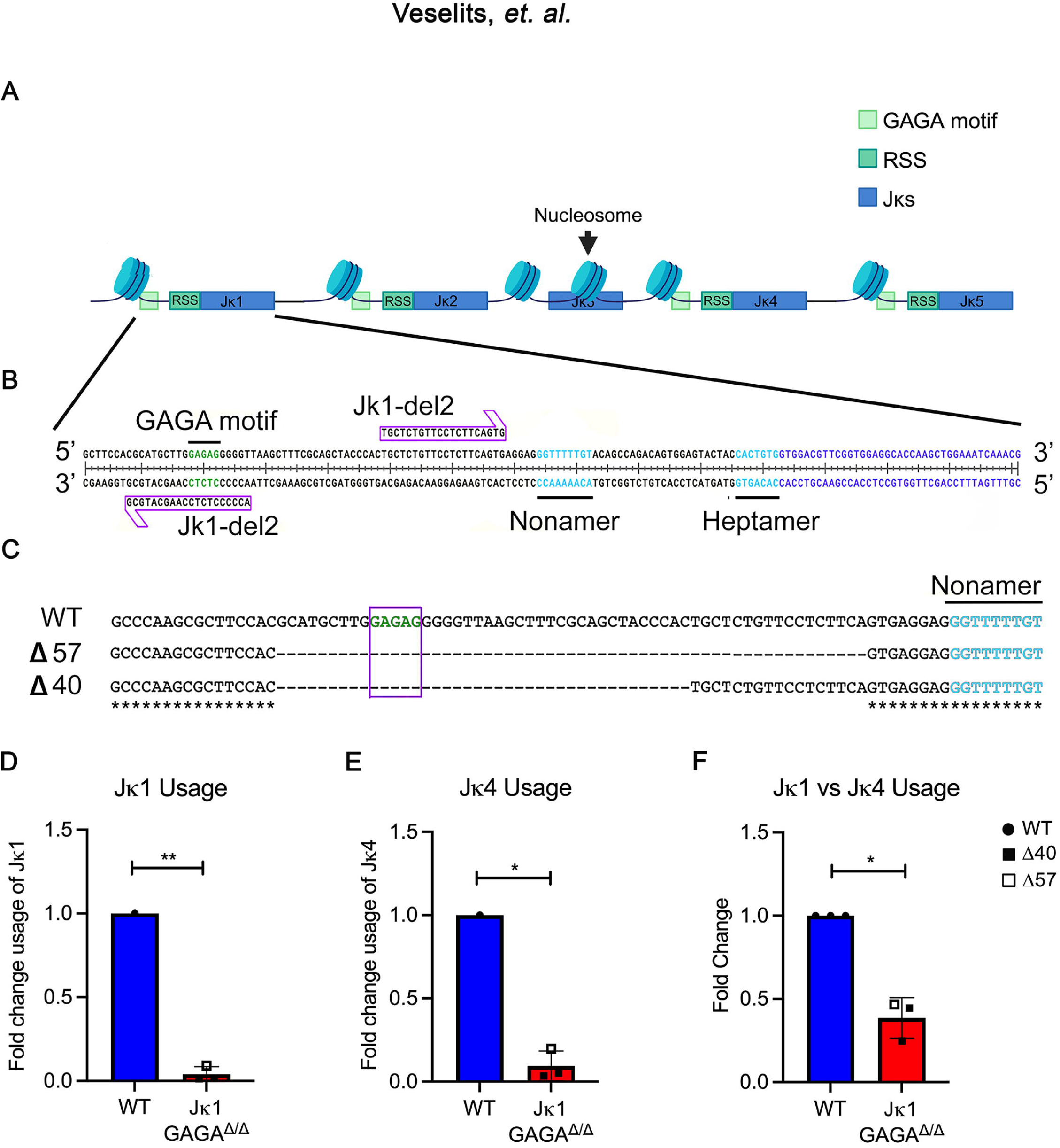
Jk1 5’ GAGA region required for *Igk* recombination. **(A)** A simplified representation of the Jk locus, which consists of 5 gene segments (blue), of which 4 functional gene segments (Jk1, Jk2, Jk4, and Jk5) are preceded by an RSS (teal) and a GAGA motif domain (green). Jk3 is a nonfunctional segment and lacks both 5’ elements. **(B)** CRISPR guides were designed to excise the smallest possible region containing the GAGA motif without disturbing the RSS. **(C)** The results of sequencing from the CRISPR-Cas9-edited mice. Two heterozygous mice with the desired deletion (40 bp and 57 bp respectively) were selected and bred to produce independent GAGA-deleted lines (“*Jk1-GAGA* ^Δ^*^40^*” and “*Jk1-GAGA* ^Δ^*^57^*”). **(D)** Quantitative RT-PCR for the Vk-Jk1 recombination product in flow-sorted small pre-B cells isolated from WT and *Jk1-GAGA* ^Δ^*^/^*^Δ^ mice (*Jk1-GAGA* ^Δ^*^40^* and *Jk1-GAGA* ^Δ^*^57^* combined in analysis, each point represents separate mouse). **(E)** Quantitative RT-PCR for the Vk-Jk4 recombination products in flow-sorted small pre-B cells isolated from WT and *Jk1-GAGA* ^Δ^*^/^*^Δ^ mice. **(F)** Quantification of relative usage of Jk1 to Jk4 in WT and *Jk1-GAGA* ^Δ^*^/^*^Δ^ mice as measured by qPCR of recombination products. (n = 3, unpaired t-test, *p <0.05, **p <0.01).

To test this prediction, we first used CRISPR-CAS9 gene editing to create a mouse model (31) in which the Jk1 upstream GAGA motif was deleted. We designed guide RNAs to cut both 5’ and 3’ to the Jk1 GAGA motif (**Figure 1B**). The guide RNAs were incubated with CAS9 protein and injected in single cell mouse embryos as a ribonucleoprotein (RNP) complex. The resulting pups displayed a variety of heterozygous mutations surrounding the Jk1 GAGA motif (**Supplementary Figure 1B**). We selected two pups with mutations that removed the GAGA sequence without disturbing the RSS or Jk1 gene body to breed to homozygosity. This resulted in two independent mouse lines in which the Jk1 5’ GAGA motif had been deleted (“*Jk1-GAGA* ^Δ^*^/^*^Δ^”, collectively), one with a 40 bp deletion (“*Jk1-GAGA* ^Δ^*^40^*”) and one with a 57 bp deletion (“*Jk1-GAGA* ^Δ^*^57^*”) (**Figure 1C**). The creation of two separate lines minimized the impact of possible confounding effects introduced by off-target mutations.

*Igk* recombination is initiated in small pre-B cells (*1, 32*). Therefore, to determine how loss of the Jk1 5’ GAGA motif region affected Jk1 usage in developing B cells, we sorted small pre-B cells (B220^+^CD19^+^CD43^-^IgM^-^FSC^low^) from wild type (WT) and *Jk1-GAGA* ^Δ^*^/^*^Δ^ mice (Supplementary Figure 2) and performed quantitative PCR (qPCR) for Vk-Jk1 recombination products (*23*). We found that in both *Jk1-GAGA* ^Δ^*^40^* and *Jk1-GAGA* ^Δ^*^57^* small pre-B cells, expression of the Vk-Jk1 recombination product was decreased nearly 40-fold (**Figure 1D**).

To assess whether recombination at other Jk segments was impacted, we performed qPCR for Vk-Jk4 recombination products. Surprisingly, Jk4 usage was also diminished in *Jk1-GAGA* ^Δ^*^/^*^Δ^ small pre-B cells, suggesting global recombination defects (**Figure 1E**). However, though the defect seemed to extend throughout the Jk locus, the effect was most profound at Jk1. The Vk-Jk1:Vk-Jk4 recombination product ratio was nearly three times lower in *Jk1-GAGA* ^Δ^*^/^*^Δ^ mice compared to WT, with both knockout murine lines providing similar results (**Figure 1F**). These data indicate that although recombination in general is decreased in the *Jk1-GAGA* ^Δ^*^/^*^Δ^ mice, the effect is most pronounced at Jk1.

### Increased Ig λ usage in Jk1-GAGA ^Δ/Δ^ mice

During immunoglobulin light chain gene recombination, developing B cells first attempt to recombine at the *Igk* locus (*1*). If this recombination is unsuccessful, impaired, or gives rise to an autoreactive BCR, B cells then attempt recombination at the *Ig*λ locus (*33, 34*). Therefore, we reasoned that if there was a meaningful recombination defect at *Igk* in the *Jk1-GAGA* ^Δ^*^/^*^Δ^ mice, there would be an increased Igλ^+^:Igk^+^ ratio in these mice relative to WT. We used flow cytometry to assess Igk vs. Igλ expression in BM immature B cells in WT and *Jk1-GAGA* ^Δ^*^/^*^Δ^ mice (**Supplementary Figure 3A**). The ratio of Igλ^+^:Igk^+^ immature B cells was indeed higher in *Jk1-GAGA* ^Δ^*^/^*^Δ^ mice (**Supplementary Figure 3B**). This reflected a decreased number of Igk^+^ immature B cells in *Jk1-GAGA* ^Δ^*^/^*^Δ^ mice while the number of Igλ^+^ cells was not significantly different. Overall, these results indicate removing the region 5’ of Jk1, that includes the GAGA motif, globally impairs *Igk* recombination.

### Deletion of Jk1 GAGA motif region alters local chromatin structure

In WT small pre-B cells, the Jk RSSs are depleted of nucleosomes, a chromatin state dictated by BRWD1 (*23*). Therefore, we next sought to determine whether deletion of the 5’ Jk1 GAGA motif domain affected Jk1 RSS nucleosome density. For this, we performed ATAC-seq (assay for transposase-accessible chromatin with sequencing) with paired-end sequencing on flow-sorted small pre-B cells from WT, *Brwd1 ^-/-^*, *Jk1-GAGA* ^Δ^*^40^*, and *Jk1-GAGA* ^Δ^*^57^* mice. Strikingly, we saw that nucleosome positioning in the Jk1 region was GAGA region-dependent (**Figure 2A**). In WT cells at the pro-B cell stage, the Jk1 gene segment and RSS were occupied by a nucleosome. Then in WT small pre-B cells, that are preparing for and undergoing *Igk* recombination, the Jk1 RSS was nucleosome-free. Instead, in small pre-B cells there was a nucleosome positioned just upstream of the Jk1 RSS (“5’ nucleosome”)(**Figure 2B**). However, in both *Jk1-GAGA* ^Δ^*^40^* and *Jk1-GAGA* ^Δ^*^57^* small pre-B cells, the 5’ region that is normally occupied by a nucleosome in WT cells remained largely clear. In contrast, in *Jk1-GAGA* ^Δ^*^/^*^Δ^ small pre-B cells, the Jk1 RSS tended to be obscured by a nucleosome (**Figure 2C)**. This defective pattern in nucleosome positioning, especially the 5’ nucleosome, was similar to that observed in *Brwd1 ^-/-^* small pre-B cells (*23*). In contrast, the nucleosome structure across the rest of the Jk gene segments was largely similar to WT cells. This is particularly evident when examining the *Jk1-GAGA* ^Δ^*^57^* mutant. From this, we conclude that without the Jk1 GAGA motif region, nucleosome structure at Jk1 is disordered.

**Figure 2.**
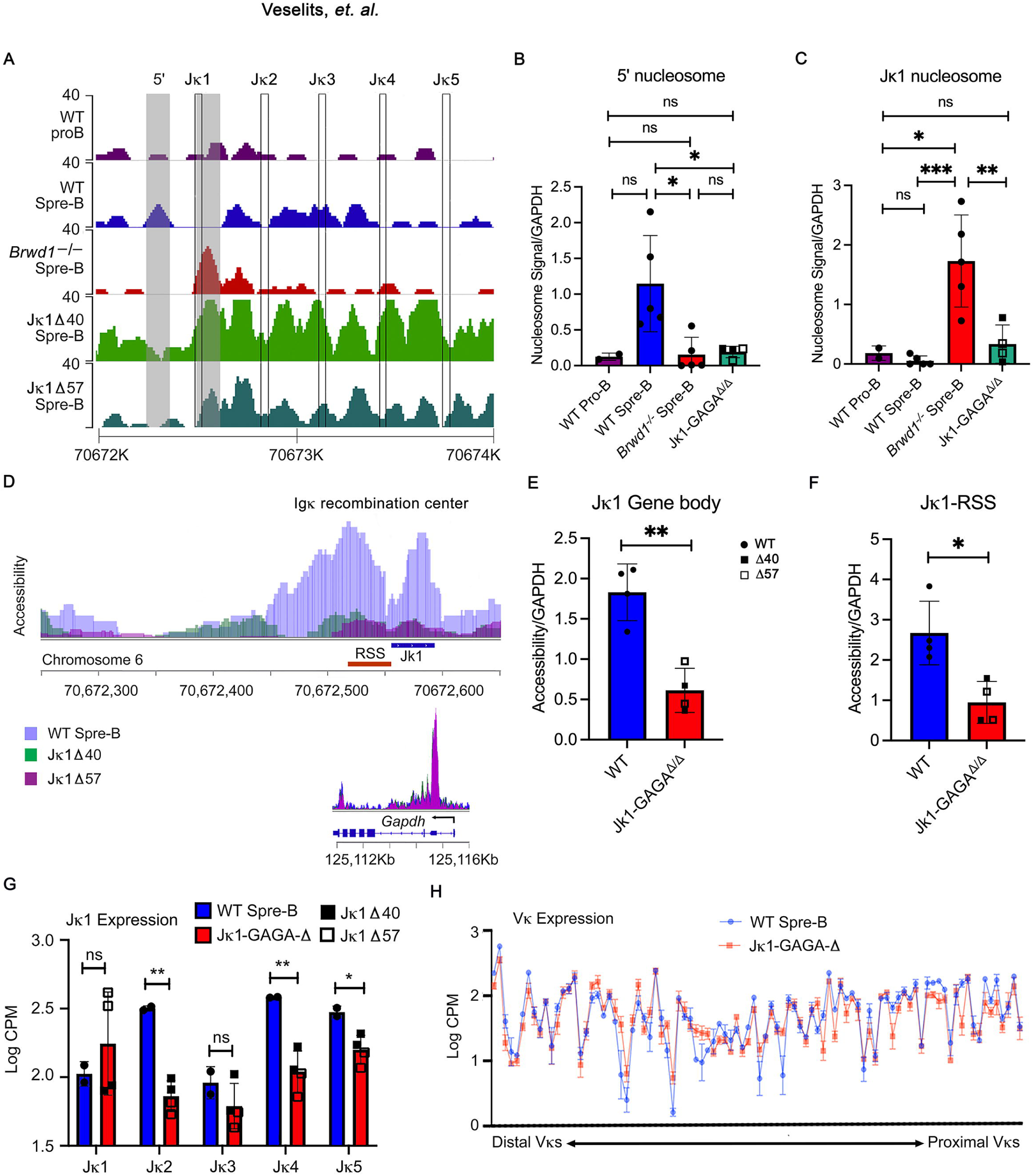
Jk1 5’ GAGA region dictates accessibility and nucleosome structure. **(A)** Nucleosome positioning at the Jk locus in WT pro B, WT small pre-B, *Brwd1 ^-/-^*small pre-B, *Jk1-GAGA* ^Δ^*^40^* small pre-B, and *Jk1-GAGA* ^Δ^*^57^* small pre-B cells. Nucleosome signal represents the difference in normalized density between the simulated signal and background data, with signal defined from read pairs with large insert sizes and background defined from read pairs with short insert sizes. Data are representative of two independent experiments. **(B)** Quantification of nucleosome signal at the indicated site 5’ of the Jk1 RSS, noted by a shaded box. **(C)** Quantification of nucleosome signal at the Jk1 gene segment plus RSS. **(D)** Accessibility at the Jk locus as measured by ATAC-seq. The y axis represents tags per million reads. Data from two independent experiments (10^5^ cells per sample). **(E)** Quantification of chromatin accessibility at the Jk1 gene body. **(F)** Quantification of chromatin accessibility at the Jk1 RSS. **(G)** Quantitation of germline Jk transcription from RNA-seq of WT vs. *Jk1-GAGA* ^Δ^*^/^*^Δ^ small pre-B cells in Log2 counts per million (CPM). **(H)** Quantification of germline Vk transcription from RNA-seq of WT vs *Jk1-GAGA* ^Δ^*^40^* and *Jk1-GAGA* ^Δ^*^57^* small pre-B cells in Log2 CPM. (unpaired t-test, *p <0.05, **p <0.01, ***p<0.001).

We also analyzed local accessibility around Jk1 in the WT, *Jk1-GAGA* ^Δ^*^40^*, and *Jk1-GAGA* ^Δ^*^57^* small pre-B cells (**Figure 2D**) compared to that at a control gene that does not undergo BRWD1-dependent remodeling, *Gapdh* . Visually, deletion of the Jk1 GAGA motif region induced large changes in local accessibility. This corresponded to decreased accessibility at the Jk1 gene body (**Figure 2E**) and a significant decrease in accessibility at the Jk1 RSS (**Figure 2F**). These data indicate that the Jk1 GAGA motif region regulates both local nucleosome structure and accessibility.

### Germline Jk1 transcription is normal in Jk1-GAGA ^Δ/Δ^ small pre-B cells

Germline transcription is induced prior to *Igk* recombination and has been tightly linked to recombination at the *Tcrd* locus (*8, 35*). However, it is unclear whether transcription reflects increased locus accessibility or if transcription plays a direct role in enhancing gene recombination. Therefore, we next assessed if the Jk1 GAGA motif region was required for Jk germline transcription. To interrogate this, we analyzed paired-end RNA-sequencing (RNA-seq) data from WT (36) as well as *Jk1-GAGA* ^Δ^*^40^* and *Jk1-GAGA* ^Δ^*^57^* small pre-B cells. To specifically analyze germline transcription and not transcription of recombination products, we removed multimapping reads from our analysis.

Throughout much of the Jk region, germline transcription of the unrecombined locus was decreased in *Jk1-GAGA* ^Δ^*^/^*^Δ^ small pre-B cells (**Figure 2G)**. The exceptions to this trend were at the Jk1 and Jk3 loci. The lack of change in expression at Jk3 is not surprising, as it is a non-functional gene segment. However, unexpectedly germline transcription at Jk1 was normal and actually trended upward. As expected, Vk germline transcription was unchanged in the *Jk1-GAGA* ^Δ^*^/^*^Δ^ mice (**Figure 2H**). These data suggest that transcription alone is insufficient to make Jk accessible for recombination.

### Jk1 GAGA motif dictates Jk1 accessibility and Vk-Jk1 recombination

In the above experiments, we deleted a small region 5’ to Jk1 containing a GAGA motif. We observed both local effects on Jk1 and a global decrease in Vk-Jk recombination. To determine if the GAGA motif contributed to local or global Jk recombination, we used CRISPR with a guide to specifically mutate the 5’ Jk1 GAGA motif (**Figure 3A**). A *Xho1* site was introduced to facilitate screening. We identified a founder in which the GAGAG motif was mutated to CTCGA (**Figure 3B**). This founder was bred to homozygosity (*Jk1-GAGA ^mut^)* and analyzed as above.

**Figure 3.**
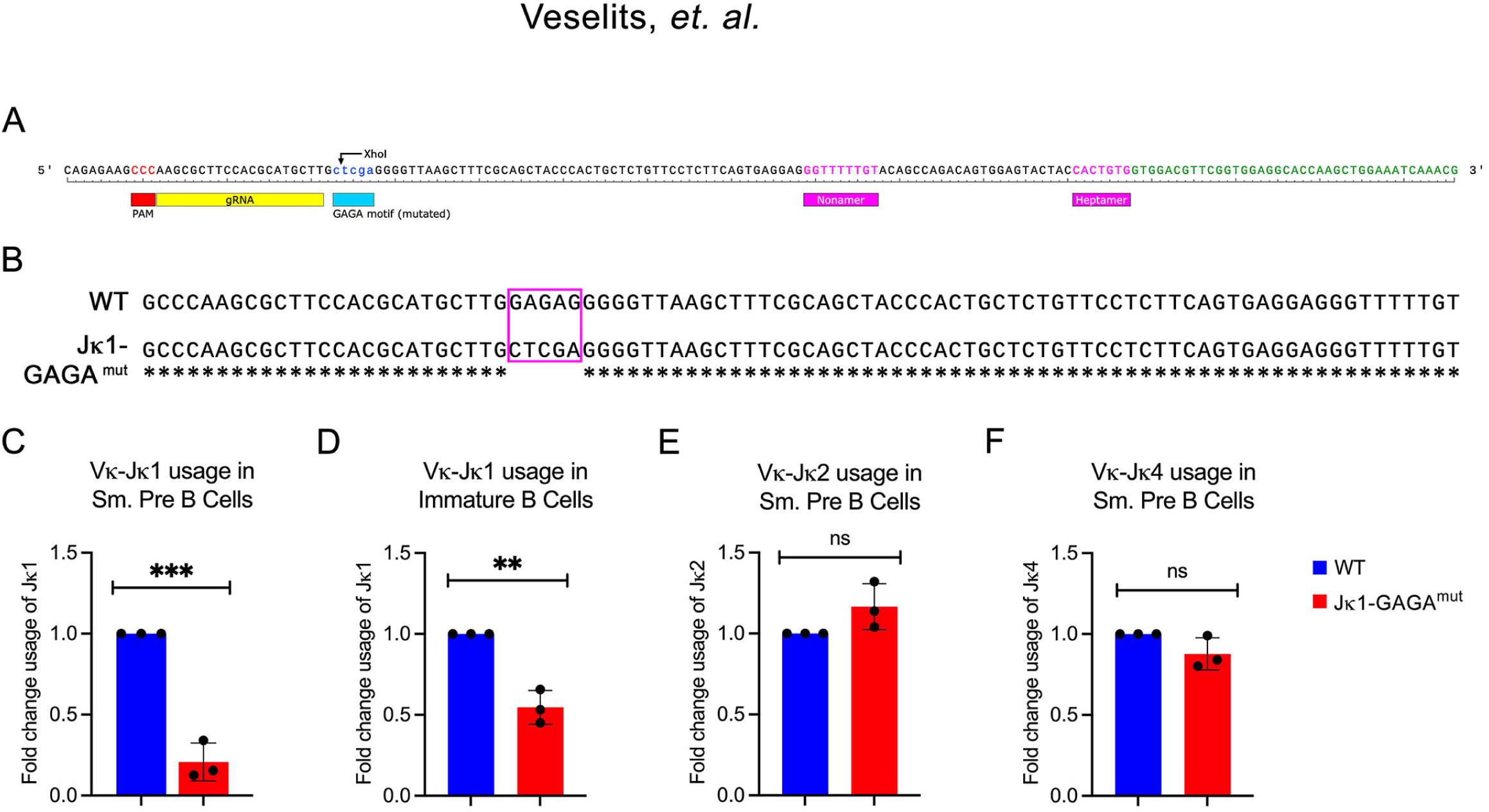
Jk1 GAGA motif required for Vk-Jk1 recombination. (**A**) Diagram of Jk1 region with guide used to mutate GAGA motif. Introduced *Xho I* sequence facilitated screening. (**B**) DNA sequence from murine founder that was then bred to homozygosity. (**C-D**) Quantitative RT-PCR for the Vk-Jk1 recombination products in flow-sorted small pre-B cells (C) and immature B cells (D) isolated from WT and *Jk1-GAGA ^mut^* small pre-B cells mice **(E-F)** Quantitative RT-PCR for Vk-Jk2 (E) and Vk-Jk4 (F) recombination products in flow-sorted small pre-B cells isolated from WT and *Jk1-GAGA ^mut^* mice. (unpaired t-test, *p <0.05, **p <0.01, ***p<0.001).

In sorted small pre-B cells, *Jk1-GAGA ^mut^* cells had severely decreased Vk-Jk1 recombination and this defect persisted into immature B cells (**Figures 3C-D**). However, Vk-Jk2 and Vk-Jk4 recombination frequencies were normal (**Figures 3E-F**). These data indicate that the Jk1 GAGA motif specifically regulates Vk-Jk1 recombination.

We next performed ATAC-seq with pair-end sequencing on the indicated flow-sorted small pre-B cell populations and assessed nucleosome placement across the Jk region (**Figure 4A**). The results were very similar to those observed in the *Jk1-GAGA* ^Δ^*^/^*^Δ^ mice with disordered nucleosome structure at Jk1 with a pronounced loss of the 5’ Jk1 nucleosome (**Figure 4B**) and a trend towards nucleosome occupancy at Jk1 gene body/RSS (**Figure 4C**).

**Figure 4.**
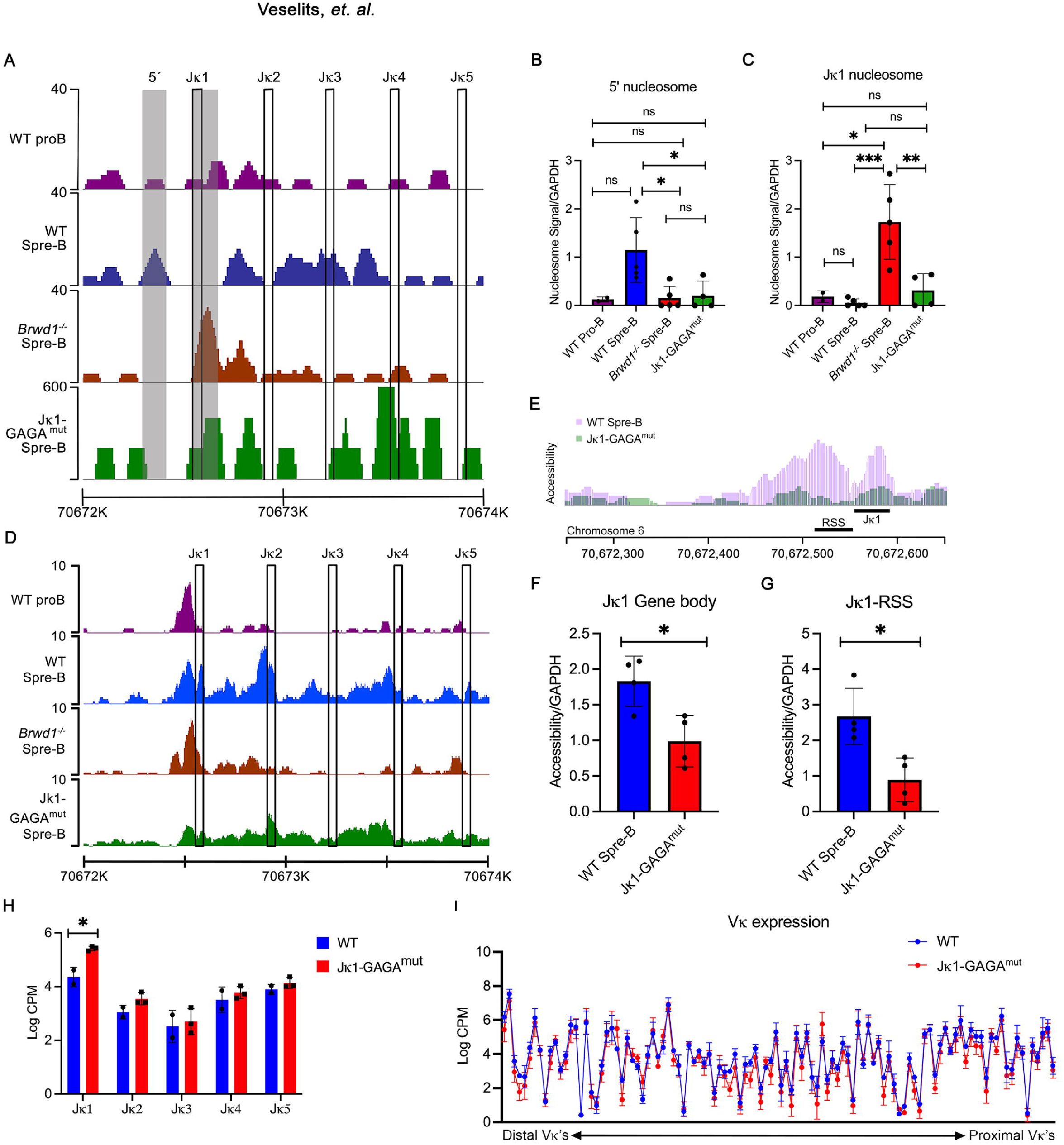
Jk1 GAGA motif required for 5’ nucleosome placement and RSS accessibly. **(A)** Nucleosome positioning at the Jk locus in WT pro B, WT small pre-B, *Brwd1 ^-/-^* small pre-B and *Jk1-GAGA ^mut^* small pre-B cells. Data are representative of two independent experiments. **(B)** Quantification of nucleosome signal at the indicated site 5’ of the Jk1 RSS, noted by a shaded box. **(C)** Quantification of nucleosome signal at the Jk1 gene segment plus RSS. **(D-E)** Examples of accessibility at the Jk locus as measured by ATAC-seq. The y axis represents tags per million reads. Data from two independent experiments. **(F)** Quantification of chromatin accessibility at the Jk1 gene body. **(G)** Quantification of chromatin accessibility at the Jk1 RSS. **(H)** Quantitation of germline Jk transcription from RNA-seq of WT vs. *Jk1-GAGA ^mut^*small pre-B cells in Log2 counts per million (CPM). **(H)** Quantification of germline Vk transcription from RNA-seq of WT vs *Jk1-GAGA ^mut^* small pre-B cells in Log2 CPM. (unpaired t-test, *p <0.05, **p <0.01, ***p<0.001).

We next analyzed our ATAC-seq data for accessibility across the Jk locus in small pre-B cells. As was observed for *Jk1-GAGA* ^Δ^ and *Brwd1 ^-/-^* cells, *Jk1-GAGA ^mut^*small pre-B cells had a global decrease in Jk1 accessibility, both at the Jk1 gene body and associated RSS (**Figures 4D-G**). Finally, analysis of Jk transcription as above in small pre-B cells revealed slightly increased Jk1 germline transcription while germline transcription of the other Jks were normal (**Figure 4H)**. As expected, Vk germline transcription was normal (**Figure 4I**). These data indicate that the Jk1 GAGA motif specifically regulates Jk1 and not other Jk genes. Sequences flanking this GAGA motif appear to have an additional role in global Jk recombination.

### Inhibition of Jk1 recombination impairs B lymphopoiesis

As the defect in *Jk1-GAGA ^mut^* mice was more specific, we further characterized these mice for developmental defects by subjecting harvested BM to flow cytometry (**Figure 5A**). Small pre-B cells, were significantly reduced in *Jk1-GAGA ^mut^*mice as compared to WT. All other BM B cell populations were normal.

**Figure 5.**
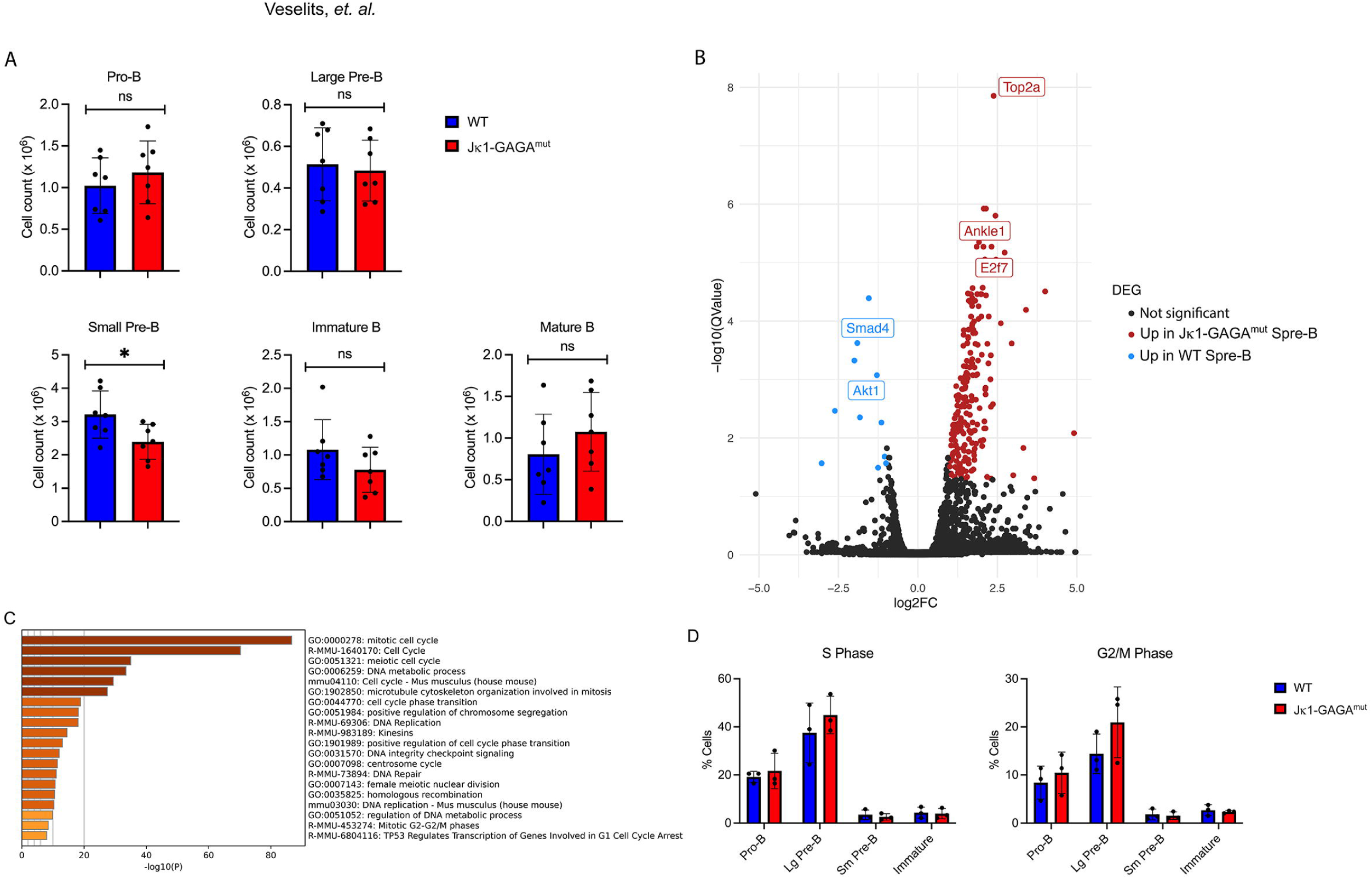
Mutation of Jk1 GAGA motif impairs B cell development. **(A)** Flow cytometric analysis of different developmental stages of B cell lymphopoiesis in the bone marrow of WT and *Jk1-GAGA ^mut^* mice. (**B)** Differential expressed genes in *Jk1-GAGA ^mut^* vs. WT small pre-B cells (n=2). Genes upregulated in *Jk1-GAGA ^mut^* cells at right with down regulated genes at left. **(C)** Gene ontology analysis of differentially expressed pathways between *Jk1-GAGA ^mut^* and WT small pre-B cells. (n = 2). **(D)** Cell cycle analysis in indicated bone marrow B cell populations. (*p < 0.05, unpaired t-test).

It was surprising that specific disruption of just Jk1 recombination would induce even a mild defect in B cell development. Therefore, we compared RNA-seq from WT and *Jk1-GAGA ^mut^* small pre-B cells. Indeed, in *GAGA ^mut^* small pre-B cells 213 genes were increased, including several genes associated with proliferation including *Top2a*, *Ankle1* and *E2f7* (**Figure 5B).** Fewer genes were downregulated. Gene ontology analysis of differentially expressed genes between *Jk1-GAGA ^mut^* and WT small pre-B cells revealed general upregulation of cell division pathways (**Figure 5C**). This was associated with a small but statistically non-significant increase of *Jk1-GAGA ^mut^* small pre-B cells in S and G2/M cell cycle phases (**Figure 5D)**. Cells must fully exit large pre-B cell proliferative programs before initiating *Igk* recombination in small pre-B cells (*1, 32*). These data suggest a mild defect in that transition. That inhibiting Jk1 recombination would impair exiting the large pre-B cell proliferative program is consistent with the known role of RAG-mediated DNA double-stranded breaks in inhibiting large pre-B cell proliferation (*37*).

### A GAGA motif is required to rescue Jk3 recombination

Jk3 is a pseudogene segment lacking both intact RSS and GAGA motifs. Remnants of both motifs are present at Jk3 suggesting evolutionarily loss of recombination competency (**Figure 6A**). Therefore, we used CRISPR-CAS9 gene editing with templates designed to repair the RSS nonamer and heptamer with or without repair of the 5’ putative GAGA motif. A *HinF1* site was introduced to facilitate screening of the RSS repair. Mice were screened by PCR and sequencing. Heterozygotes were bred to homozygosity to generate Jk3^GAGA/RSS^ and Jk3^RSS^ mice (**Figures 6B** and **C** respectively).

**Figure 6.**
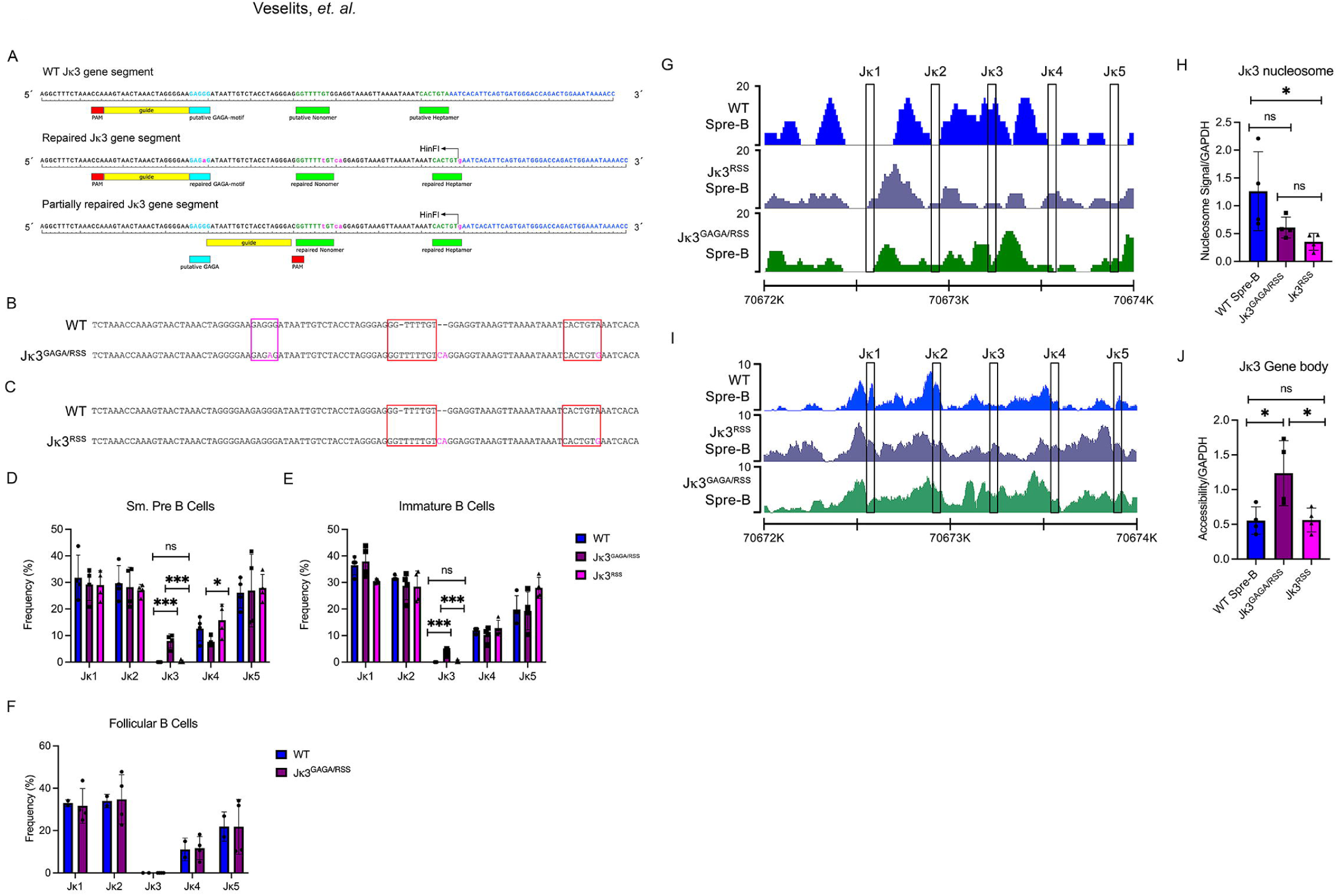
Both RSS and GAGA motifs required to rescue Vk-Jk3 recombination. (**A**) Diagram of Jk3 region with partially repaired (repaired RSS) and fully repaired (RSS and GAGA motif) sequences shown. Shown are the guides used. See methods for repair templates. Introduced *HinFI* sequence facilitated screening. (**B-C**) DNA sequence from murine founders that were then bred to homozygosity. (**D-F**) Vk-Jk gene products were amplified by PCR and individual clones sequenced. Shown are the frequencies of indicated Vk-Jk gene products from WT, *Jk3 ^GAGA/RSS^* and *Jk3 ^RSS^* mice from flow-sorted small pre-B (D), immature B (E) and follicular B (F) cells. **(G)** Nucleosome positioning at the Jk locus in small pre-B cells from WT, *Jk3 ^GAGA/RSS^* and *Jk3 ^RSS^* mice. Data are representative of two independent experiments. **(H)** Quantification of nucleosome signal over Jk3 gene body and RSS. **(I)** Example of accessibility at the Jk locus in indicated mouse strains as measured by ATAC-seq. The y axis represents tags per million reads. Data representative of two independent experiments. **(J)** Quantification of chromatin accessibility at the Jk3 gene body and RSS. (unpaired t-test, *p <0.05, **p <0.01, ***p<0.001).

No validated PCR primers are available for Jk3. Therefore, we sorted small pre-B single cells from WT, Jk3^GAGA/RSS^ and Jk3^RSS^ mice, isolated mRNA and from the resulting cDNA performed PCR with degenerate Vk primers and a Ck primer. Individual PCR products were cloned and sequenced. All three mouse lines demonstrated similar frequencies of Vk-Jk products containing Jk1, Jk2 and Jk5. Jk3^GAGA/RSS^ had a small decrease in Jk4 containing recombination products (**Figure 6D**). Remarkably, 7.5% (27/358 sequences) of Jk3^GAGA/RSS^ small pre-B cells contained Vk-Jk3 expressed sequences. In contrast, only 1 of 400 sequenced PCR products (0.25%) from single Jk3^RSS^ small pre-B cells contained a Vk-Jk3 product. Therefore, both a GAGA motif and a RSS are necessary and sufficient for recombination to Jk gene segments in small pre-B cells.

We next assessed Vk-Jk3 recombination frequencies in BM immature B cells and splenic follicular B cells (**Figures 6E and 6F**). In immature B cells, 3.9% percent of sequences contained Vk-Jk3 products (15/384). In contrast, we detected no Vk-Jk3 expressed gene products (0/200) in follicular B cells. These data are consistent with negative selection of B cells expressing immunoglobulin receptors encoded by Vk-Jk3 and suggest that the rescued Jk3 confers autoreactivity.

We then used ATAC-seq data to compare nucleosome positioning at Jk3 in WT small pre-B, Jk3^GAGA/RSS^ small pre-B and Jk3^RSS^ small pre-B cells (**Figure 6G**). In WT small pre-B cells, Jk3 is enriched for nucleosomes including 5’ to the Jk3 gene segment (**Figure 6H)**. The presence of a 5’ nucleosome in WT cells obscured potential contributions of the restored GAGA motif at this position. However, distinct nucleosome structure over the Jk3 RSS and gene body was diminished in both Jk3^GAGA/RSS^ small pre-B and Jk3^RSS^ small pre-B cells. However, these changes were only statistically significant for Jk3^RSS^.

We then used the ATAC-seq data to examine Jk accessibility (**Figure 6I**). In WT small pre-B cells Jk3 RSS gene segment accessibility was low and this increased in Jk3^GAGA/RSS^ small pre-B cells (**Figure 6J**). Interestingly, accessibility was not conferred by addition of only an RSS. These observations suggest that the primary function of a GAGA motif at Jk3 is to enhance accessibility across the RSS/Jk3 region.

To begin to understand if GAGA motifs were used at other recombination centers, we examined 5’ nucleotide sequences at the murine IgHJ (JH) and TCRαJ (Jα) gene segments (**Supplementary Figure 4**). Indeed, GAGAv motifs were found 5’ to all JH segments, which, like Jk, contain 23bp RSSs. At these sites, the GAGA motifs tended to be found around 320 bp (range: 273-361 bp) 5’ to the RSS nonamer. We also found GAGA motifs 5’ to all Jα gene segments examined (15/15). However, the spacing at Jα genes between GAGA and the corresponding RSS was variable. The Jα cluster contains 12 bp RSSs. These data suggest that GAGA motifs might be important at recombination centers other than that at Jk.

### Increased nucleosome occupancy at cryptic 23 RSSs

Our data support a model in which one of the activities of GAGA motifs is to make Jk RSSs accessible in preparation for recombination. This model is consistent with *in vitro* data indicating that nucleosome occupancy protects RSSs from RAG-mediated cleavage (*20*). These findings led us to consider whether similar mechanisms might repress DNA cleavage at the many cryptic RSSs scattered throughout the genome.

Broadly, the genome is organized into compartments (*38, 39*). Compartment A contains actively transcribed genes and open chromatin while Compartment B consists of repressed genes packaged into heterochromatin. Chromatin compartments are cell lineage and state specific. It has previously been shown that genome-wide there has been evolutionary selection against cRSSs (*13*). As the risk of RAG-mediated cleavage should be higher in open chromatin, we first examined if cRSSs were preferentially depleted in Compartment A.

We defined A and B compartment boundaries using Hi-C data obtained from WT C57BL/6 small pre-B cells (*40*). Next, we compiled an annotation of all 12 and 23 cRSSs passing an established threshold throughout the entire mouse genome using the RSS Information Content (RIC) algorithm (*41–43*). We were able to identify approximately 3.55 x 10^6^ cRSSs. We observed that the density of cRSSs in Compartment A appeared lower than in Compartment B (**Figure 7A)**. Using the density of cRSSs throughout the entire genome, we were able to calculate predicted numbers of cRSSs in Compartment A and Compartment B if cRSSs were to be distributed entirely randomly throughout the genome (**Figure 7B**). This revealed that cRSSs are relatively depleted from Compartment A and enriched in Compartment B. These data suggest that the evolutionary pressure against cRSSs is preferentially confined to those that lie in the open chromatin of developing B cells.

**Figure 7.**
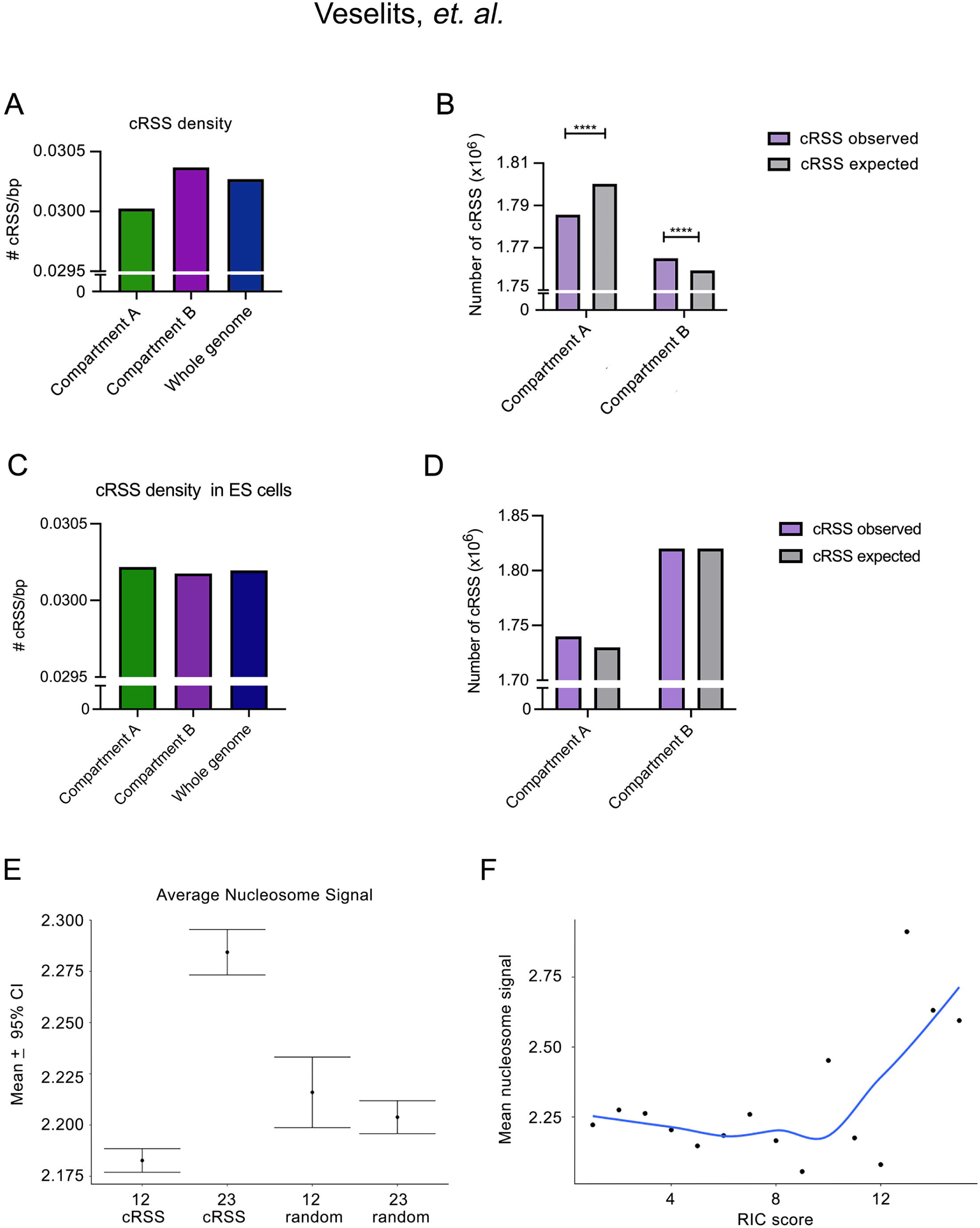
Cryptic RSSs are depleted in active chromatin and bind nucleosomes. **(A)** Cryptic RSS density in indicated genome Compartments in WT small pre-B cells. (**B**) Total number of observed cRSSs per small pre-B cell Compartment versus the number of cRSSs expected by random distribution of cRSSs throughout the genome (p < 0.0001, Chi-squared test). (**C**) Cryptic RSS density in indicated genome Compartments in WT embryonic stem cells. (**D**) Total number of observed cRSSs per embryonic stem cell Compartment versus the number of cRSSs expected by random distribution of cRSSs throughout the genome. **(E)** Mean nucleosome signal in WT small pre-B cells for 12 and 23 cRSSs compared to randomly generated sequences of the same length. Plotted as mean +/- a 95% confidence interval. **(F)** Mean nucleosome signal in small pre-B cells vs RIC Score.

We next asked if selection against cRSSs in compartment A was specific to B cells. Therefore, we examined cRSS density in Compartment A and B of C57BL/6 embryonic stem (ES) cells (**Figure 7C**). As can be seen, cRSS densities in Compartment A and B were similar with a slight increase in Compartment A. Likewise, the number of expected versus observed cRSSs in both Compartment A and B were similar with a slight, but not statistically significant, increase in observed cRSS in Compartment A (**Figure 7D**). These data suggest that evolutionary selection against cRSSs is determined by the chromatin accessibility of those cells undergoing RAG-mediated recombination.

Next, we determined whether cRSSs are more likely to be occupied by nucleosomes and are thus relatively shielded from errant recombination. We first used our previously published ATAC-seq data from WT small pre-B cells to calculate nucleosome density across the whole genome (*36*). We then mapped annotated cRSSs in Compartment A and compared this to one million randomly chosen Compartment A genomic regions corresponding to the length of either a 12 or a 23 RSS. We used the bedtools bigWigAverageOverBed tool to determine mean nucleosome occupancy (**Figure 7E**). The 23 cRSSs had a significantly higher mean nucleosome signal than either the randomly chosen 12 or 23 regions. However, interestingly, the 12 cRSSs had a significantly lower mean nucleosome signal. This suggested that 12 cRSSs and 23 cRSSs are epigenetically regulated in different manners.

Finally, we wanted to determine if nucleosome occupancy correlated with the strength of the RIC score. The algorithm that determines the RIC score ranks a sequence on how closely it resembles a canonical RSS, with higher scores being closer to canonical. Most RSSs are low scoring and only poorly resemble a canonical RSS. We bucketed and ranked the RIC scores from weakest (RIC score = 1) to strongest (RIC score = 15) and plotted the mean nucleosome score. This analysis revealed that as an RSS becomes stronger, the mean nucleosome signal increased (**Figure 7F**). Together, these data suggest that in open chromatin cRSSs, particularly strong 23bp cRSSs, are preferentially bound to nucleosomes *in vivo* .

## Discussion

Herein, we demonstrate that the canonical Jk RSSs are not sufficient to ensure efficient RAG-mediated genomic cleavage and recombination. Rather, a local 5’ GAGA motif is essential for efficient recombination to the Jk RSSs and for dictating local nucleosome structure and genomic accessibility. While traditionally the focus has been on the RAG cleavage site encoded by the RSS, we now demonstrate the importance of associated DNA GAGA motifs which ensure the Jk epigenetic state required for Vk-Jk recombination.

Our data support a model in which the presence of a 5’ GAGA motif ensures the epigenetic landscape necessary for efficient recombination in two primary ways. First, the GAGA motif ensures that the RSS is accessible, which *in vitro* studies have shown is required for permitting RAG1-mediated RSS cleavage (*20*). Second, the GAGA motif directs positioning of a nucleosome 5’ to the RSS, which is predicted to enable precise recruitment of RAG2 (*17–19*). Therefore, our findings suggest an expanded definition of the minimal motif necessary RAG-mediated cleavage at a recombination center.

Our experiments demonstrated that a 5’ GAGA motif was required for efficient recombination to both Jk1 and a reconstituted Jk3. The effect of each GAGA motif, on both nucleosome placement and recombination, was primarily restricted to the proximate Jk gene segment. However, larger gene deletions 5’ to the Jk1 RSS, that deleted the GAGA motif and surrounding sequences, induced global changes in the Jk region with generally diminished Vk-Jk recombination. These data suggest that beyond the Jk1 GAGA motif, there are additional elements 5’ to Jk1 important for Jk recombination.

There are several plausible models by which Jk segments might be chosen. The first is the stochastic model, by which Jk segments are chosen for recombination purely at random. Another model is the sequential model, by which recombination is first tried at Jk1, and if that fails, recombination can be tried sequentially 5’ to 3’, from Jk2 to Jk5. Our experiments in which we manipulate GAGAs at both Jk1 and Jk3 are consistent with stochastic choice of Jk gene segments for recombination. However, the mild developmental defect seen in *Jk1-GAGA ^mut^* mice suggest that recombination to Jk1 is attempted early.

Data from the Jk1 and Jk3 mutant mice show a clear dependency of nucleosome positioning on GAGA motifs. It has been previously shown that BRWD1 is recruited to the Jk locus and plays an important role in both clearing local RSSs of nucleosomes and 5’ nucleosome placement (*23*). Furthermore, nucleosome positioning in Jk1 GAGA-mutant small pre-B cell cells resembled that of the *Brwd1 ^-/^*^-^ cells at the Jk1 locus.

BRWD1 is recruited genome-wide to specific epigenetic landscapes which do or do not contain extended GAGA motifs. Therefore, GAGA motifs are not required for BRWD1 genome recruitment. However, BRWD1-dependent nucleosome remodeling occurs only when GAGA motifs are present (*23*). Thus, the overall picture is most consistent with GAGA Jk motifs serving as a guide for nucleosome positioning by BRWD1. This is consistent with the role GAGA motifs play in chromatin remodeling in *Drosophila* (*28*).

Asymmetry was observed between 12bp and 23bp cRSSs. Cryptic 23 RSSs were much more likely to be occupied by nucleosomes, suggesting that they have an intrinsic proclivity to bind nucleosomes *in vivo* . This was especially true for strong cRSSs. This is predicted to preferentially protect 23 cRSSs from aberrant cleavage and thus confer protection for genomic integrity. In contrast, we did not observe this putative genomic protective mechanism at 12 cRSSs. A higher intrinsic binding propensity could facilitate nucleosome loading, which is likely required for RAG2 recruitment. In this model, the Jk 23 RSSs would bind a nucleosome which will later be slid 5’ by a GAGA motif-guided, BRWD1-dependent mechanism in preparation for recombination (*23*). Our data suggest the mechanism at J 12bp RSSs, which occur at T cell recombination centers, might be different.

In addition to revealing a specific mechanism protecting 23bp cRSSs, our analysis revealed a general selection against cRSSs in the open chromatin of developing lymphocytes. It was previously known that there was a strong evolutionary pressure against cRSSs genome-wide (*13*). However, we have further demonstrated that the transcriptionally active Compartment A, in which *Igk* lies in developing B cells, is relatively cRSS depleted. In contrast, the inactive and more compacted, and thus intrinsically safer, Compartment B shows a relative increase in cRSS density. No such selection against ES cell Compartment A cRSSs was observed. These findings suggest that genomic risk from off-target RAG-mediated cleavage in lymphocytes drove genome-wide evolution.

Our findings reveal that an extended recombination motif at Jk, including both of the RSS and 5’ GAGA motifs, is critical for gene recombination. While the RSS encodes the necessary cleavage site for the RAG recombinase, the 5’ GAGA motif dictates the local chromatin architecture required for efficient recombination. These findings likely have implications for other recombination events involving RAG recruitment. Indeed, we found GAGA motifs 5’ to both JH and Jα RSSs suggesting a general requirement for antigen receptor gene recombination. Therefore, the GAGA motif-dependent mechanism we described in small pre-B cells, might ensure the fidelity and contextual appropriateness of RAG-mediated gene recombination in other lymphocyte populations.

## Materials and Methods

### Mice

Wild-type (C57BL/6), *Jk1-GAGA* ^Δ^*^40^* (C57BL/6), *Jk1-GAGA* ^Δ^*^57^* (C57BL/6), *Jk1-GAGA ^mut^*(C57BL/6)*, Jk3 ^GAGA/RSS^* (C57BL/6)*, Jk3 ^RSS^* (C57BL/6) and *Brwd1* ^-/-^ (C57BL/6-C3HeB/FeJ) mice were housed in the animal facility of the University of Chicago. Male and female mice were used at 6-12 weeks of age in accordance with the Institutional Animal Care and Use Committee of the University of Chicago.

### CRISPR-Cas9 Gene Editing

CRISPR-Cas9 RNAs (crRNAs) were designed to flank the regions targeted for deletion using the crRNA design resource hosted by the Feng Zhang lab and CHOPCHOP (Supplementary Table 1) (*44, 45*). crRNAs were optimized for high efficiency and low off-target scores and were synthesized by Integrated DNA Technologies (IDT) as Alt-R CRISPR-Cas9 crRNA oligos. To produce the guide RNA (gRNA), crRNA and tracrRNA (IDT) were resuspended at 1 µg/µl in injection buffer (1 mM Tris-HCl, pH 7.5, 0.1 mM EDTA), mixed at a 1:2 ratio by mass (5 µg crRNA and 10 µg tracrRNA), and annealed in a thermocycler (95 °C for 5 min, ramp down to 25 °C at 5 °C/min).

To produce the active ribonucleoprotein (RNP) mixture, a 100 µl solution was prepared containing the gRNA and Alt-R S.p. Cas9 Nuclease (IDT). For the *Jk1-GAGA* ^Δ^*^/^*^Δ^ mice, the RNP mix was prepared with 25 ng/µl of each gRNA and 100 ng/µl of Cas9. For the *Jk1-GAGA ^mut^, Jk3 ^GAGA/RSS^ and Jk3 ^RSS^* mice, 300 ng/µl of gRNA and 300 ng/µl of Cas9 was used as well as 100 ng/µl of Cas9 mRNA. The RNP mix was incubated at room temperature for 15 min after which 200 ng/μl of ssODN repair template was added. The RNP mix was then centrifuged at 13,000 RPM at room temperature to remove solid particles, and the top 80 µl was used. Injections were performed by the University of Chicago Transgenics Core. One cell fertilized embryos were injected with the RNP mix and implanted into pseudopregnant mice.

Due to the heterogeneous nature of CRISPR-Cas9 gene editing, both alleles were screened for edits separately in the F0 generation. The region surrounding the targeted DNA was amplified by PCR using primers listed in Supplementary Table 1, and products were cloned into a TA vector. Genotyping vectors were transformed and a minimum of 8 individual colonies were sequenced.

### ATAC-seq

ATAC-seq was performed with 1.2 x 10^5^ FACS-sorted small pre-B and immature B cells as previously described (*23, 36*). Briefly, cells were washed with PBS and lysed with ATAC lysis buffer (10 mM Tris-HCl, pH 7.4, 10 mM NaCl, 3 mM MgCl2, 0.1% IGEPAL CA-630). Nuclei were incubated with the transposase tagmentation enzyme (Illumina). Library fragments were amplified using the Nextera Indexing kit (Illumina) and NEBNext PCR master mix (New England BioLabs) for 10-12 cycles and were purified with the QIAquick PCR Purification Kit (Qiagen). Libraries were size-selected with the E-Gel SizeSelect gel system (Life Technologies) in the range of 150-650 bp. We quantified the size-selected libraries with an Agilent Bioanalyzer and via qPCR in triplicate using the KAPA Library Quantification Kit on the Life Technologies Step One System. Libraries were sequenced on the Illumina HiSeq2000.

ATAC-seq data was analyzed as previously described (*23, 36*). For comparative analysis of chromatin accessibility between samples, we used the bedtools bigWigAverageOverBed, with the open chromatin bigWig file and a bed file containing annotated regions of interest as inputs. Samples were normalized to accessibility at *Gapdh* . Similarly, for comparative analysis of nucleosome occupancy between samples, we instead used the nucleosome signal bigWig file.

### Quantitative PCR, RNA-seq, and analysis

Total cellular RNA was isolated by resuspending cells in 500 μl of TRIzol for 5 min at room temperature (Invitrogen), then 100 µl of chloroform was added and incubated at room temperature for 2-3 min. Samples were centrifuged for 15 min at 12,000 g at 4 °C, and 0.25 ml of isopropanol was added to the aqueous phase. Samples were centrifuged for 10 min at 12,000 g at 4°C. The RNA pellet was resuspended in 1 ml of 75% ethanol and centrifuged for 5 min at 7500 x g at 4 °C. The supernatant was discarded, and the pellet was resuspended in 20 μl of DNase/RNase-free water.

For quantitative PCR, total cellular RNA was reverse transcribed with SuperScript III reverse transcriptase (Invitrogen). A total volume of 25 μl containing 1 μl cDNA template, 0.5 μM of each primer (Supplementary Table 1) and SYBR Green PCRMaster Mix (Applied Biosystems) was analyzed in triplicate. Gene expression was analyzed with an ABI PRISM 7300 Sequence Detector and ABI Prism Sequence Detection Software version 1.9.1 (Applied Biosystems). Normalized results were calculated using the ΔΔCt method (Expression Fold Change = 2^-((KO_TEST_- KO_HOUSEKEEPING_) - (WT_TEST_- WT_HOUSEKEEPING_)) using *B2m* as the housekeeping gene.

For RNA-seq, RNA libraries were prepared using the standard Illumina library protocol (Kit, RS-122-2101 TruSeq Stranded mRNA LT-SetA) before sequencing on the Illumina HiSeq2500. Raw reads were aligned to reference genome mm9 in a splice-aware manner using STAR (*46*). Gene expression was quantified using FeatureCounts against UCSC genes, with Ensembl IG genes from mm10 converted to mm9 coordinates with UCSC liftOver (*47*).

Differential expression statistics (fold-change and p value) were computed using edgeR on raw expression counts obtained from quantification (*48*). Pairwise comparisons were computed using exactTest, and multigroup comparisons using the generalized linear modeling capability in edgeR. In all cases, p values were adjusted for multiple testing using the FDR Benjamini-Hochberg correction. Significant genes were determined based on an FDR threshold of 0.05 in the multigroup comparison. Metascape was used for pathway analyses (*49*).

### PCR analysis of *Igk* rearrangements

Quantification of Jk usage by PCR was performed by sequencing the products of *Igk* rearrangements. Degenerate Vk and Ck primers were used along with 2 μl of cDNA in a 25 μl reaction using Platinum Taq DNA polymerase (ThermoFisher). Two μl of the PCR product was cloned into the pCRII-TOPO TA vector (ThermoFisher) and transformed into DH5α cells (ThermoFisher). DNA was extracted from resulting colonies with the QIAprep Spin Miniprep Kit (Qiagen) and sequenced. Unique sequences were analyzed for Jk usage by alignment to the mouse Jk (Supplementary Table 1) and Vk sequences using the NCBI BLAST alignment tool (https://blast.ncbi.nlm.nih.gov/).

### Flow cytometry and flow activated cell sorting (FACS)

Bone marrow was extracted from the hind leg bones of mice, suspended in RPMI media with 10% vol/vol FBS, and passed through a 70 μm filter. Red blood cells were lysed with ACK lysis buffer (Lonza). Cells were then washed and resuspended in staining buffer (PBS with 3% vol/vol FBS) and blocked with 2.5 μl of Fc block (BD Pharmingen) for 30 min. Cells were stained with anti-B220-PerCP-Cy5.5 (RA3-6B2), anti-CD19-APC-Cy7 (1D3), anti-CD43-PE (S7), anti-IgM-APC (II/41), anti-Igk-BV510 (187.1), anti-Igλ-FITC (R26-46). Small pre-B cells (B220^+^CD43^-^IgM^-^FSC^low^) and immature B cells (B220^+^CD43^-^IgM^+^) were isolated by cell sorting with a FACSAriaII (BD). Flow cytometric analysis was done with FlowJo (BD).

### Hi-C compartment analysis

Hi-C data on murine pre-B cells has previously been published (*50*). The genome was divided into A and B chromatin compartments based on the correlation matrix (”runHiCpca.pl”). A and B compartments were based on the eigenvector sign, with compartment A defined as a positive eigenvector value and compartment B defined as a negative eigenvector value (*51–54*).

### cRSS identification and analysis

The online Recombination Signal Sequences Site (https://www.itb.cnr.it/rss/) was used to identify cRSSs (*43*). A RIC score threshold of -38.81 was used to identify 12 cRSSs and a threshold of -58.45 was used to identify 23 cRSSs across each chromosome (*55*). The cRSS start site, end site, chromosome, RIC score, and a unique identifier were combined in a bed file. To exclude *Igk* and *Ig*λ RSSs, a bed file of *Igk* and *Ig*λ RSSs was used, and overlapping sites were removed using the bedtools intersect tool with option - v. The final bed file had 3,559,481 total cRSSs.

To produce a non-cRSS outgroup for comparison, the bedtools random tool was used to produce 1,000,000 random tracks of length 27 (equivalent to 12 cRSS) and 1,000,000 random tracks of length 38 (equivalent to 23 cRSS) with the seed set to 1. To remove any sequences from the random tracks with a cRSS, the random tracks were run through bedtools overlap -v with the cRSS bed files. The final random tracks contained 923,381 random 27 bp sequences and 911,267 random 38 bp sequences.

To calculate nucleosome average and sum density over all cRSSs and random sequences, the bedtools bigWigAverageOverBed tool was used with bigwig files of nucleosome occupancy produced by the DANPOS tool as described previously (*23*). The average nucleosome occupancy score was 95.

To correlate the strength of the RIC score and the mean nucleosome occupancy, the 12 and 23 cRSSs were first separated. cRSSs were broken into 15 bucketed RIC score ranges from weakest (RIC score = 1) to strongest (RIC score = 15). For each range, the mean nucleosome occupancy was calculated. The 12 and 23 cRSS data was reaggregated and a linear regression was performed.

To analyze cRSSs in A and B compartments, the compartment annotations were separated into two bed files for compartment A and B. To make bed files with only cRSSs in compartments A or B, the bedtools intersect tool was used with the cRSS bed file with the option -wa. 1,793,006 cRSSs were in compartment A and 1,766,138 cRSSs were in compartment B. To calculate the density of cRSSs in each compartment, the total number of bps comprising each compartment was calculated. Compartment A was 59,471,819 bps and compartment B was 58,121,675 bps.

## Acknowledgements

The authors have no conflicting financial interests.

---

**Supplementary Table 1.**
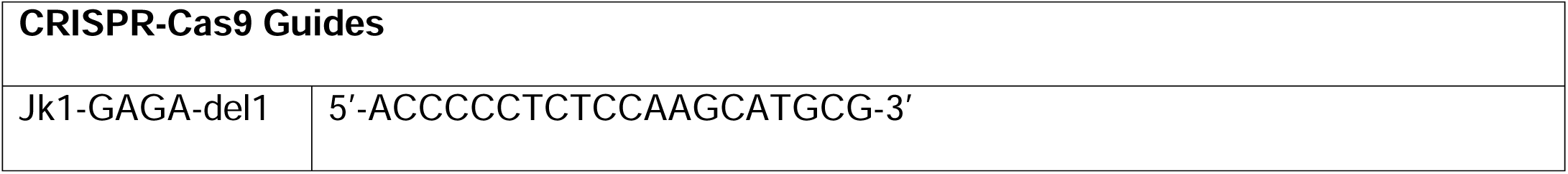
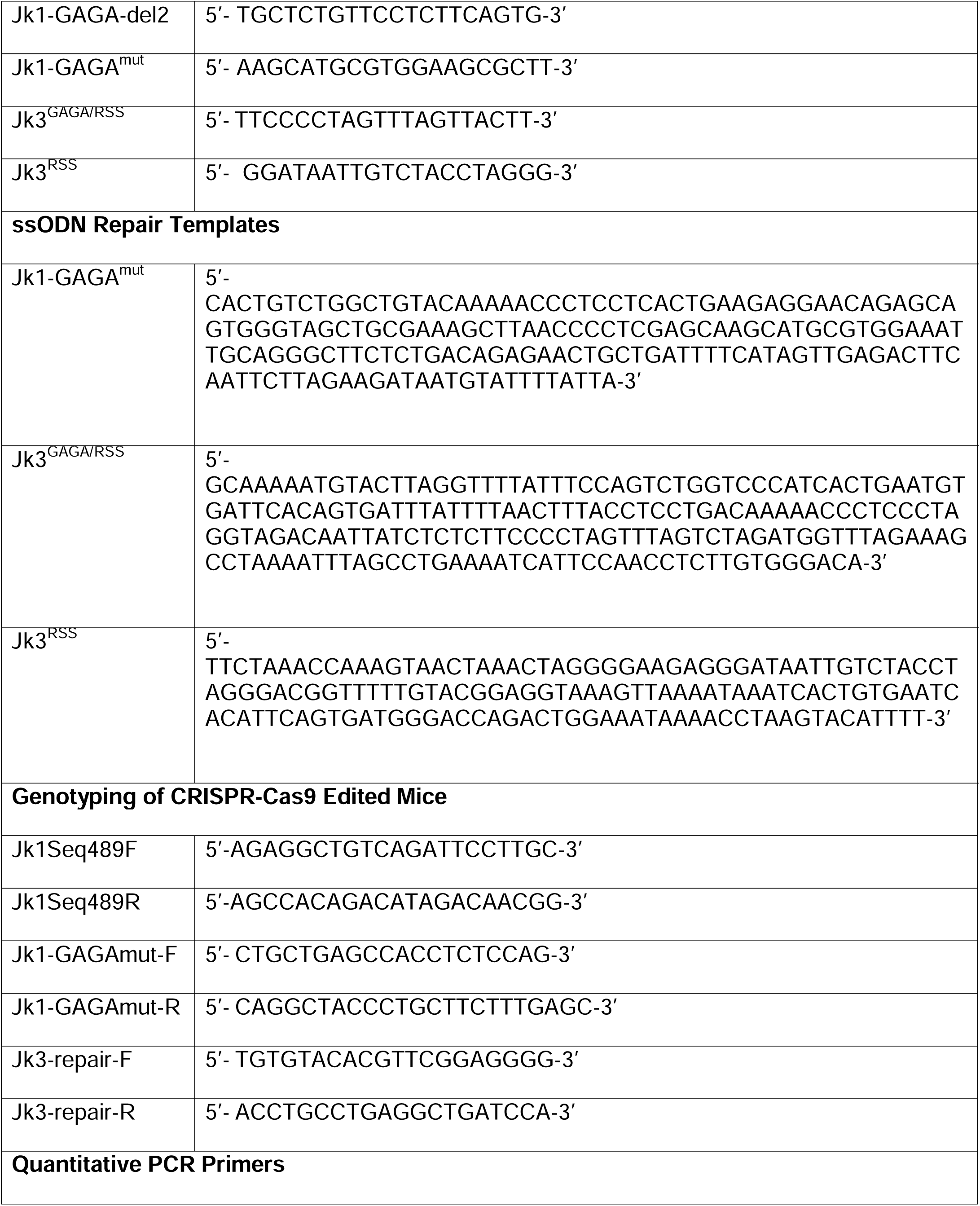
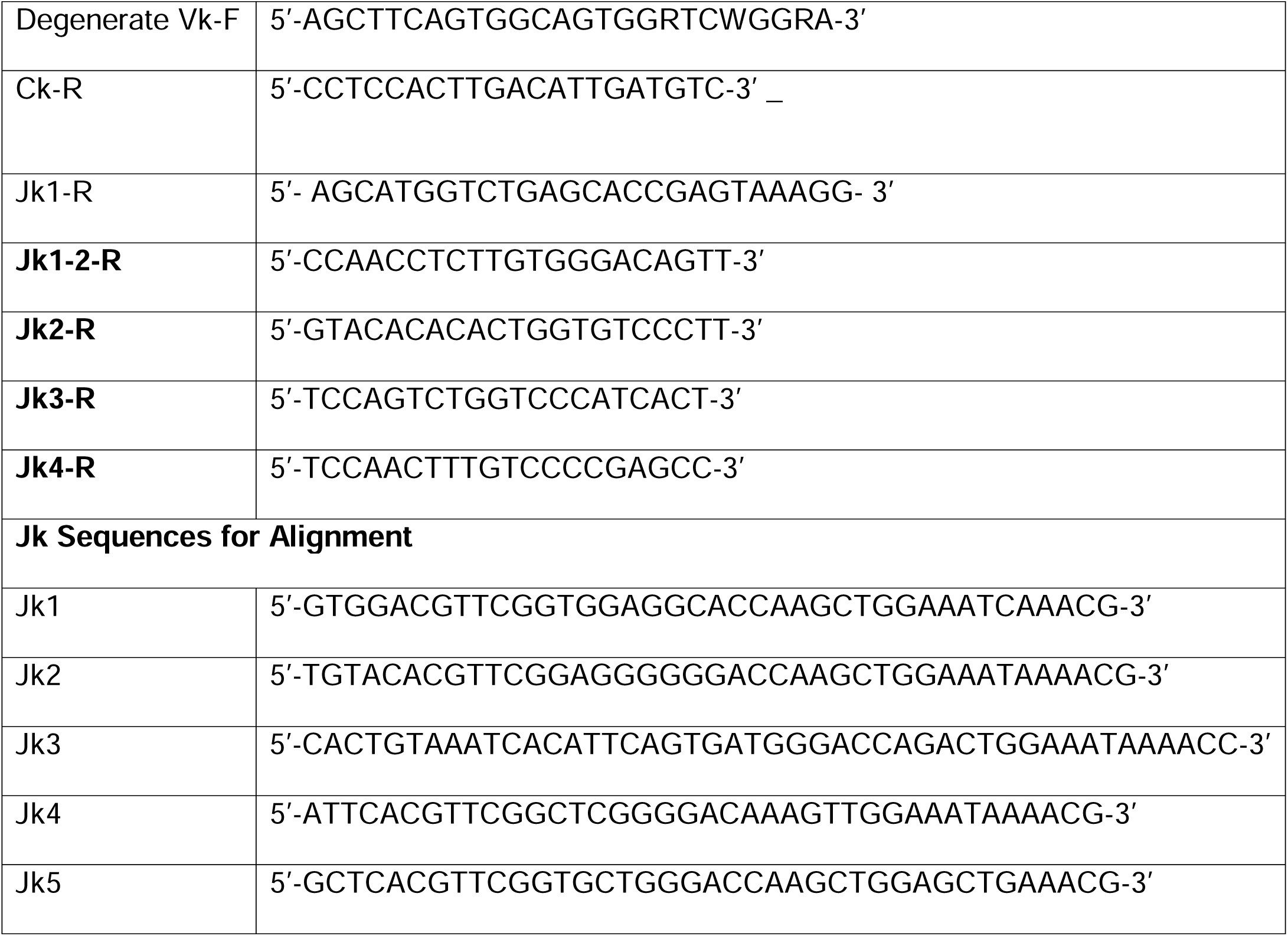
List of gRNAs and primers.

## Supplementary Figure Legends

**Supplementary Figure S1. Deletion of conserved Jk1 5’ GAGA motif**

**(A)** A GAGAG DNA pentamer (“GAGA motif”) located ∼80 bp or ∼350 bp upstream of the Jk1 gene segment is conserved across queried vertebrate species. **(B)** An alignment of all mutations found in the F0 CRISPR-Cas9-edited mice. Two heterozygous mice with the desired mutation were selected for breeding to homozygosity (Figure 1).

**Supplementary Figure S2. Flow cytometry of bone marrow B cell populations.** Flow cytometric analysis of indicated bone marrow B cell populations in WT vs *Jk1-GAGA* ^Δ^*^/^*^Δ^ mice.

**Supplementary Figure 3. Preserved Ig**λ **expression in *GAGA***^Δ^***^/^***^Δ^ **mice.** (**A**) Example of flow cytometry for Igλ and Igκ surface expression in immature B cells of indicated genotypes. (**B**) Ratios and absolute cell numbers per mouse of Igκ and Igλ expressing cells in immature B cells of the indicated genotypes. (unpaired t-test, *p <0.05).

**Supplementary Figure 4. GAGA motifs at other recombination centers.** Aligned sequences showing GAGA motifs (red), RSSs (blue) and gene body (green) for IgHJ genes (upper sequences) and TCRαJ segments (middle and lower sequences).

