## Supplementary figures and images for "Jk DNA GAGA MOTIFS ARE REQUIRED FOR LOCAL NUCLEOSOME REMODELING AND Vk-Jk RECOMBINATION"

### Supplemental Figure 1

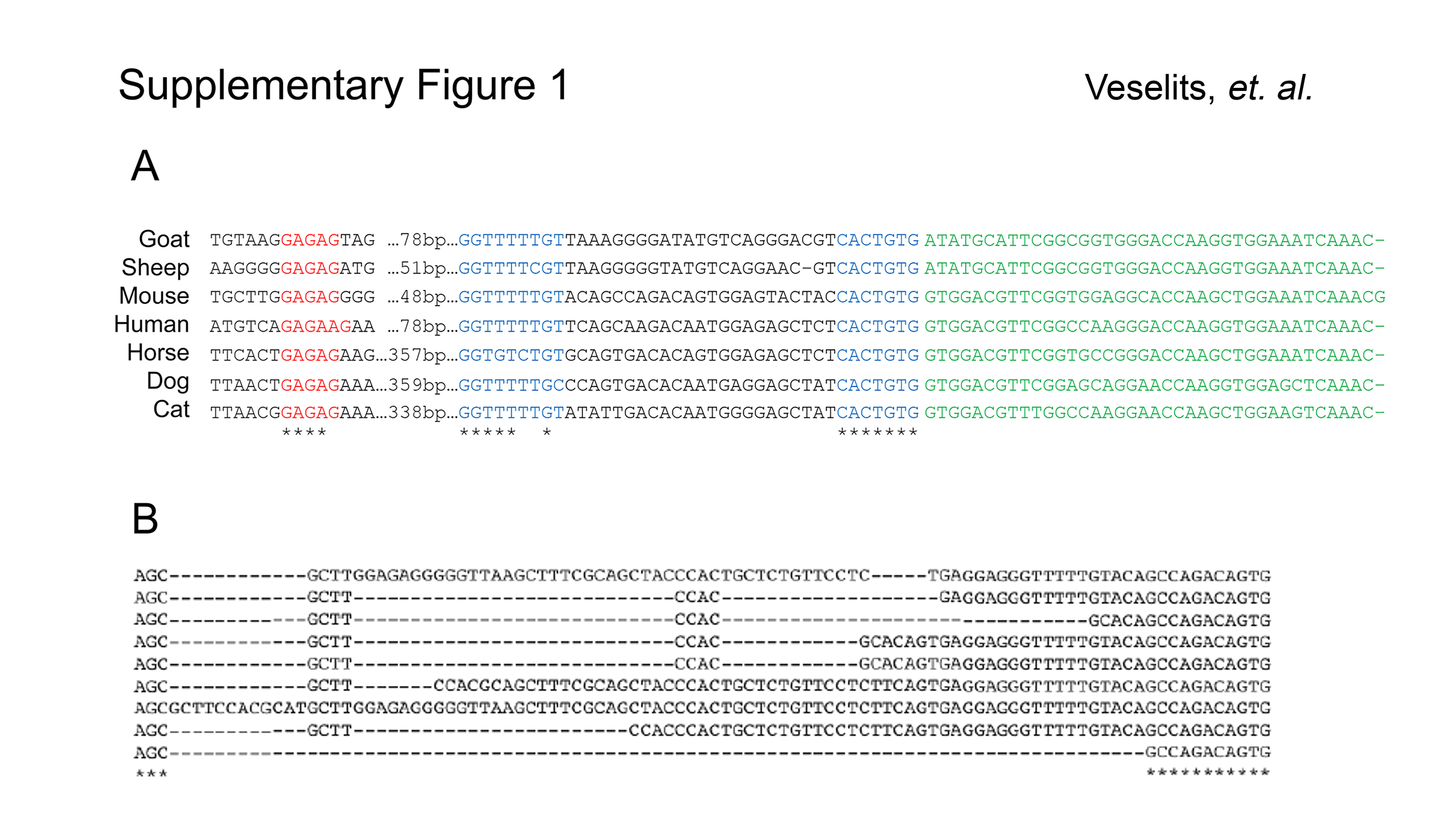

### Supplemental Figure 2

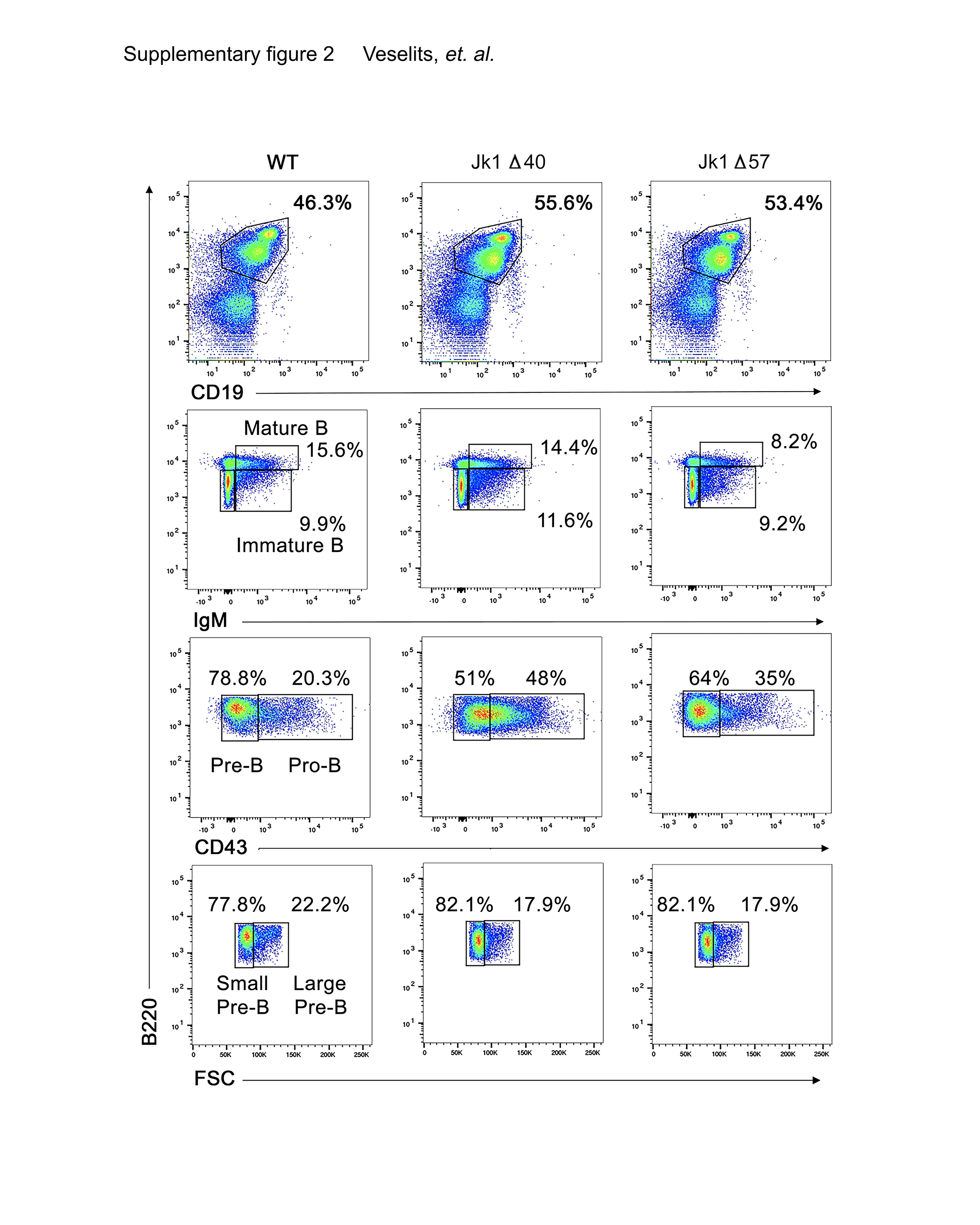

### Supplemental Figure 3

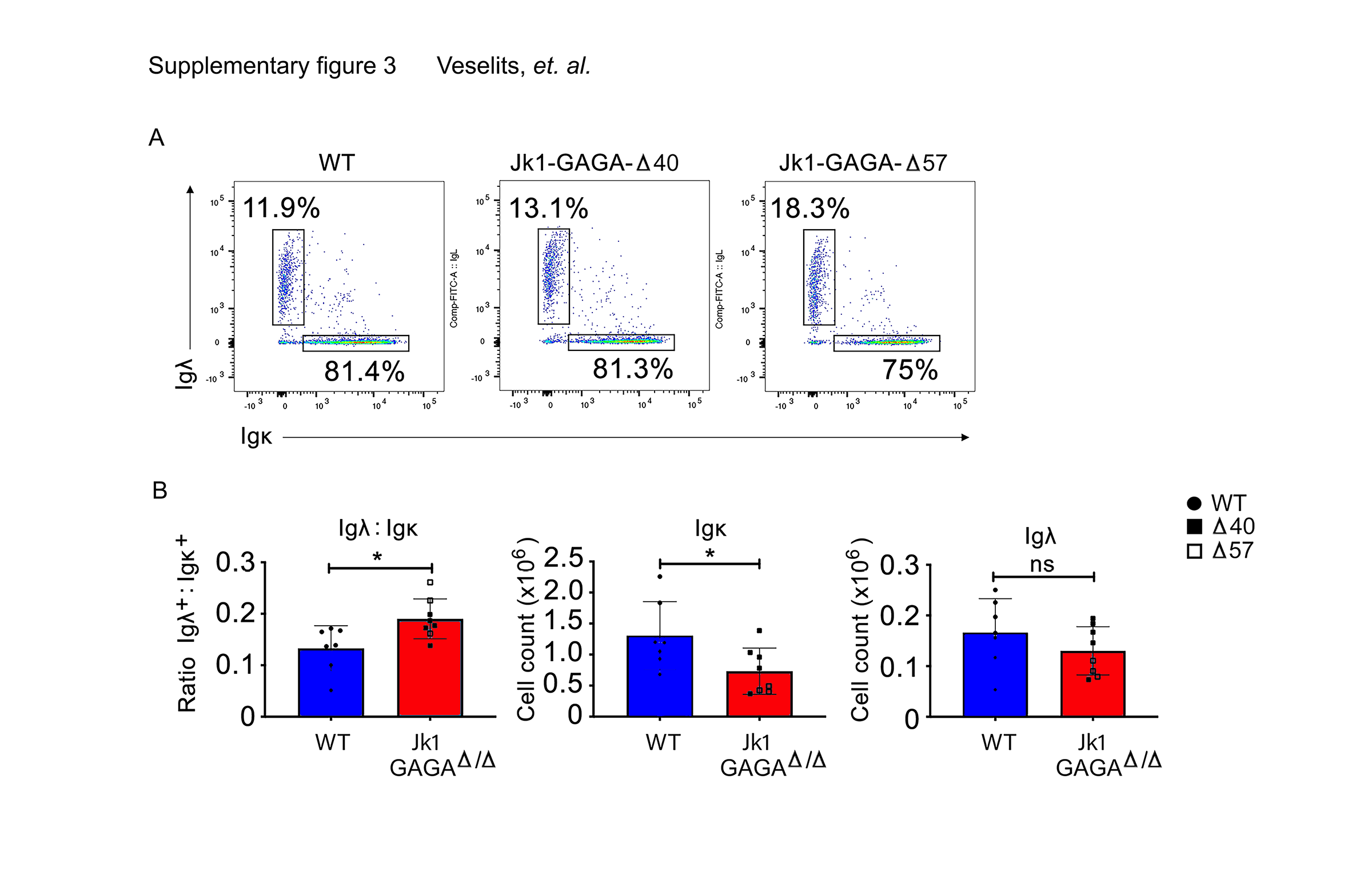

### Supplemental Figure 4

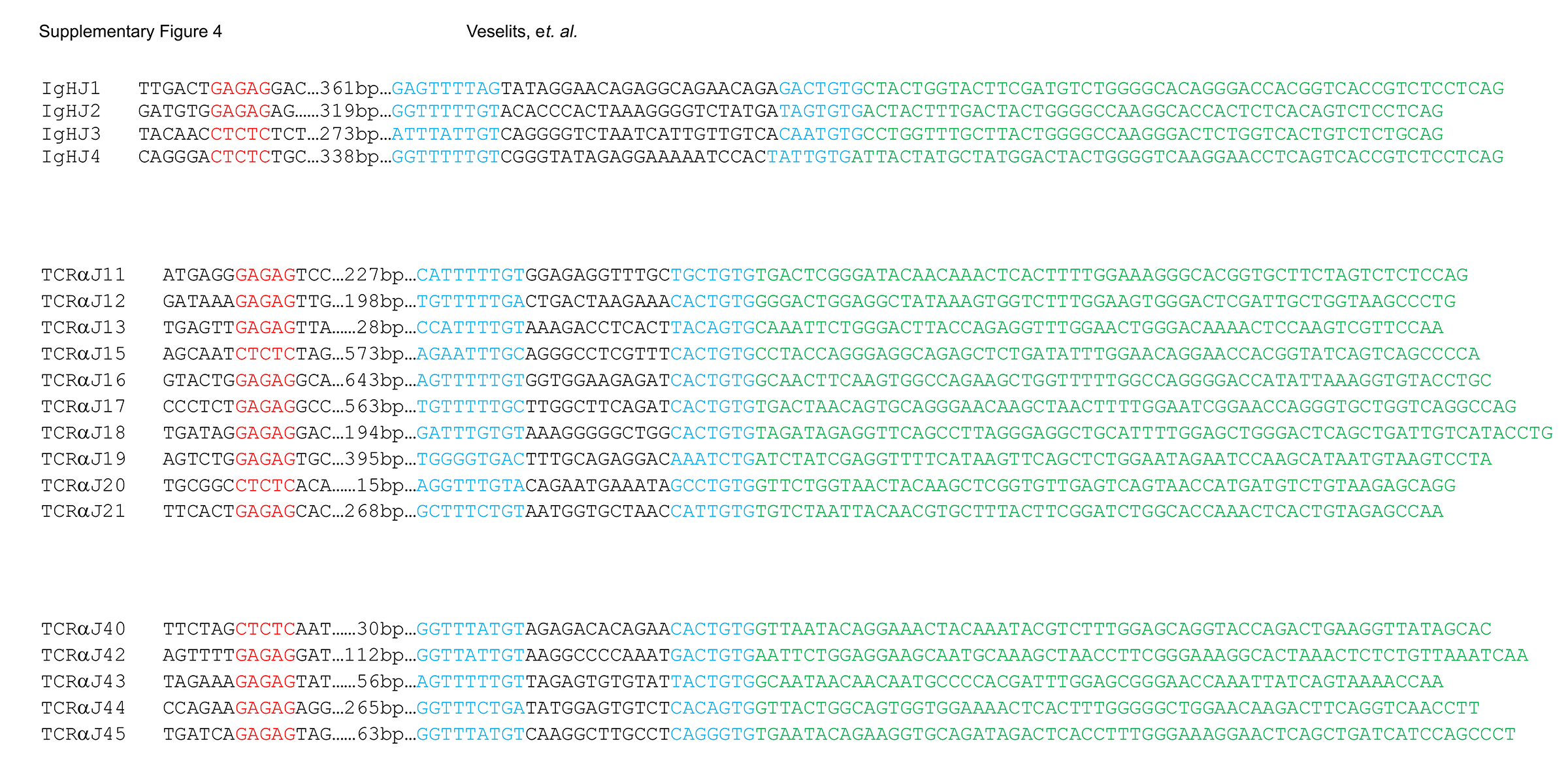
